# Characterization of primary structure of the major silk gene, *h-fibroin*, across caddisfly suborders (Trichoptera)

**DOI:** 10.1101/2022.08.29.505681

**Authors:** Jacqueline Heckenhauer, Russell J. Stewart, Blanca Ríos-Touma, Ashlyn Powell, Tshering Dorji, Paul B. Frandsen, Steffen U. Pauls

## Abstract

**Background:** Larvae of Trichoptera produce silk to build various underwater structures allowing them to exploit a wide range of aquatic environments. The *heavy chain fibroin* (*h-fibroin*) gene encodes the primary protein component of their silk. Studies on this long (>20 kbp) and highly repetitive gene have been limited by difficulties in its sequence assembly. Recently, high-quality long-read sequencing techniques have been successfully applied to obtain the full-length *h-fibroin* sequence.

**Results:** We used three new and five previously published genomes to identify eight full-length h-fibroin gene and protein sequences of h-fibroin across the order Trichoptera covering various endpoints of the diversity of ecological silk use in caddisflies. We analyze these together with four existing high-quality h-fibroin sequences. Across the order, we observed conserved patterns in h-fibroin (high similarity of amino(n)-/carboxyl(c)-termini, presence of characteristic repeating structural modules). However, the sequence, number, and arrangement of these repeating modules varied across clades with increasing structural complexity of h-fibroin in fixed retreat and tube-case builders compared to cocoon-builders. We also found a higher percentage of proline in fixed-retreat makers.

**Conclusion:** This study provides characterizations of the primary structure of h-fibroin from a diverse set of caddisflies. The interplay of conserved termini and basic motif structure with high variation in repeating modules as well as the variation in the percentage of proline might be linked to differences in mechanical properties (i.e., tensile strength, toughness) related to the different silk usage. This sets a starting point for future studies to screen and correlate amino acid motifs and other sequence features with quantifiable silk properties.

## Background

Insects use silk for a variety of purposes and most research has focused on terrestrial Lepidoptera (moths and butterflies), especially on the commercially important silkworm *Bombyx mori*. Less well-studied are aquatic insects that use silk. These include the most speciose aquatic insect order, Trichoptera (caddisflies), which exhibit diverse silk usage strategies. Similar to their sister order Lepidoptera, several caddisfly species construct cocoons for metamorphosis in the final larval instar. In earlier larval stages, they produce a diverse array of underwater structures which is reflected in their phylogeny. Trichoptera is divided into two suborders, which are distinguished by differences in morphology, habitat, and use of silk [1]: Annulipalpia (fixed retreat makers) and Integripalpia (cocoon and tube case makers). Further differentiation in silk use also occurs at the subtordinal and superfamily level, and in some cases even among congeneric species.

Caddisfly silk adheres to various substrates underwater and has high tensile strength, extensibility, and toughness [2]. Because of these unique qualities it is of interest as a model for biomimetic adhesives. Their diverse net- and case-making behavior allow caddisflies to exploit a range of ecological niches, which raises the question of how the properties of underwater silk evolved in Trichoptera. To begin to answer this question, and to gauge the potential of caddisfly silk in material sciences, a comprehensive study of the primary molecular structure of the major silk protein, h-fibroin, is necessary.

The silk fiber in Trichoptera consists of two filaments derived from a pair of labial glands (see also [3]). The fiber core is assembled from a large (200-500 kDa) heavy-chain fibroin (h-fibroin) and the smaller (~25 kDa) light chain fibroin (l-fibroin) protein. The h-fibroin is the major silk protein by size and mass. It consists of non-repetitive amino(n)- and carboxyl(c)-terminal domains flanking a central region, composed of repeated structural modules [4,5]. These consist of repeating (SX)_*n*_E motifs in which the S (serine) is often phosphorylated [6–8] and where X is primarily an amino acid with hydrophobic or aromatic side chains and sometimes arginine, E is glutamic acid, and n is 3-5. The (SX)_*n*_ E motifs are separated by glycine-rich regions of variable length [4,9,6].

The molecular structure of full-length *h-fibroin* sequences has not been studied extensively, due to challenges in the assembly of the *h-fibroin* caused by its length (>20 kilobase pairs [kbp]) and highly repetitive regions [10–12]. Only partial *h-fibroin* sequences of Trichoptera were derived from sequencing the ends of cDNAs [4,13,6]. The combination of long- and short-read sequencing approaches resulted in the assembly of two full-length *h-fibroin* sequences [10,14]. However, the *h-fibroin* could not be assembled in more than 20 species with these sequencing techniques [15,16]. Recently, new genomic long-read sequencing techniques with low sequencing error rates as well as newly developed bioinformatic tools allowed for full-length assembly of the *h-fibroin* gene sequences of four Trichoptera species (two retreat makers [12,17]; one retreat- and one cocoon-maker, [17].

In this study, we increased the number of high-quality full-length Trichoptera *h-fibroin* sequences from four to twelve. Specifically, we identified complete protein-coding *h-fibroin* gene sequences from five publicly available high-quality genomes (four case- and one retreat maker) and used PacBio HiFi to sequence and assemble the full-length *h-fibroin* sequence of three additional caddisfly species, one for each major silk usage strategy (fixed retreat, cocoon and tube case makers). We then characterized and compared the primary structure of these h-fibroins.

## Data description

### New genomic resources

Aquatic insects have been neglected with respect to genome sequencing efforts [18]. This lack of well-resolved genome assemblies has hindered the progress in understanding the genomic basis of aquatic insect traits, such as silk. Recent studies showed that new high-quality, long-read sequencing techniques allow for assembly of the repetitive >20-kbp *h-fibroin* gene [11,12,17]. We therefore used previously published and novel genomic high-quality resources produced in this study to identify and compare the gene structure of full-length caddisfly *h-fibroin* sequences.

To complement the public data that we analyzed, we extracted and sequenced genomic DNA using PacBio HiFi sequencing for three species from each major type of silk usage in Trichoptera: *L. lineaticorne* (retreat-/capture net maker, ~24.5x sequencing coverage), *Himalopsyche tibetana* (cocoon maker, ~23.3x) and *M. longulum* (tube case maker, ~15x). We assembled the HiFi reads and assessed assembly quality in four ways: (1) we checked for contiguity by calculating assembly statistics (e.g. N50) using Quast v5.0.2, (2) assed completeness by screening for single-copy orthologs with BUSCO v5.2.2, (3) calculated the back-mapping rate of the raw reads to the assembly, and (4) searched for potential contaminations using BlobTools v1.1.1 [19]. Our *de novo* assemblies of *L. lineaticorne* and *H. tibetana* rank among the highest quality assemblies for Trichoptera with respect to gene completeness (i.e., more than 96% complete BUSCOs, assembly lengths congruent with estimated genome size, table 1) and contiguity (e.g., contig N50: ~29 Mbp in *H. tibetana*; number of contigs: 65 in *L. lineaticorne*, table 1). Re-mapping of the raw reads to the assembly revealed that 98.2% (*H. tibetana*) and 99.94% (*L.lineaticorne*) could be unambiguously placed with expected coverage distribution per position (supplementary figures 1 & 3). BlobTools detected no contaminations (supplementary figures 2 & 4). Due to low sequencing coverage of *M. longulum*, the assembly was of poor contiguity. Details on assemblies are given in supplementary note 1.

**Table 1:** Comparison of currently published high quality long-read-based genome assemblies of caddisflies. **\* this study. BUSCO: % of complete BUSCOs is given based on** BUSCO 5.2.2 [42] using the endopterygota_odb10 dataset. C: complete, S: single, D: duplicated, F: fragmented, M: Missing

| Species | Suborder (silk usage) | Accession number | Assembly length (bp) | Contig N50 (contig / scaffold) | No. of contigs / scaffolds | % BUSCO (n=2124) |
| --- | --- | --- | --- | --- | --- | --- |
| *Atopsyche davidsoni* [54] | Basal Integripalpia (cocoon) | GCA_022113835.1 | 370,818,532 | 14,095,054 / n.a. | 80 / n.a. | C:96.9%[S:96.1%,D:0.8%],F:1.8%,M:1.3% |
| *Himalopsyche tibetana* * | Basal Integripalpia (cocoon) | JAPJYX000000000 | 691,323,649 | 28,889,006 / n.a. | 282 / n.a. | C:96.5%[S:95.6%,D:0.9%],F:2.4%,M:1.1% |
| *Eubasilissa regina* [11] | Integripalpia (tube case) | GCA_022840565.1 | 917,621,729 | 32,427,664 / n.a. | 123 / n.a. | C:95.5%[S:94.8%,D:0.7%],F:3.0%,M:1.5% |
| *Glyptotaelius pellucidus* (DtoL) | Integripalpia (tube case) | GCA_936435175.1 | 1,037,123,706 | 8,185,058 / 36,814,344 | 285 / 57 | C:90.3%[S:89.5%,D:0.8%],F:6.8%,M:2.9% |
| *Hesperophylax magnus* [12] | Integripalpia (tube case) | JAIUSX000000000 | 1215205050 | 11,205,906 / n.a. | 980 / n.a. | C:91.4%[S:89.0%,D:2.4%],F:6.0%,M:2.6% |
| *Limnephilus lunatus* (DtoL) | Integripalpia (tube case) | GCA_917563855.2 | 1,269,651,477 | 18,993,099 / 95,392,806 | 139 / 39 | C:89.7%[S:88.9%,D:0.8%],F:7.4%,M:2.9% |
| *Limnephilus marmoratus* (DtoL) | Integripalpia (tube case) | GCA_917880885.1 | 1,629,971,709 | 8,018,677 / 56,174,236 | 395 / 68 | C:90.4%[S:89.3%,D:1.1%],F:6.7%,M:2.9% |
| *Limnephilus rhombicus* (DtoL) | Integripalpia (tube case) | GCA_929108145.2 | 1,578,808,083 | 10,796,652 / 54,234,467 | 272 / 62 | C:89.8%[S:88.6%,D:1.2%],F:7.1%,M:3.1% |
| *Leptonema lineaticorne* * | Annulipalpia (retreat) | GCA_024500535.1 | 273,010,349 | 13,827,090 / n.a. | 65 / n.a. | C:96.1%[S:95.3%,D:0.8%],F:2.3%,M:1.6% |
| *Cheumatopsyche charites* [48] | Annulipalpia (retreat) | GCA_024721215.1 | 223,232,897 | 2,851,765 / 13,966,006 | 207 / 68 | C:96.4%[S:95.9%,D:0.5%],F:1.9%,M:1.7% |
| *Arctopsyche grandis* [17] | Annulipalpia (retreat) | n.a., see Frandsen et al, 2022 | 485,663,687 | 6,470,670 / n.a. | 676 / n.a. | 97.3%[S:93.5%,D:3.8%],F:1.6%,M:1.1% |

### Identification, annotation and comparison of h-fibroin sequences

We identified *h-fibroin* genes in the two newly and five previously published genome assemblies (table 2, supplementary table 1), annotated these and checked for their quality by searching their sequences for stop codons, frame shift errors, and spurious introns. In addition, we confirmed that there is a signal peptide at the beginning of each gene. Although the assembly quality of *M. longulum* was low, we were still able to assemble and annotate the *h-fibroin* gene, rendering the species’ sequences useful for comparison of *h-fibroin* genes.

**Table 2:**
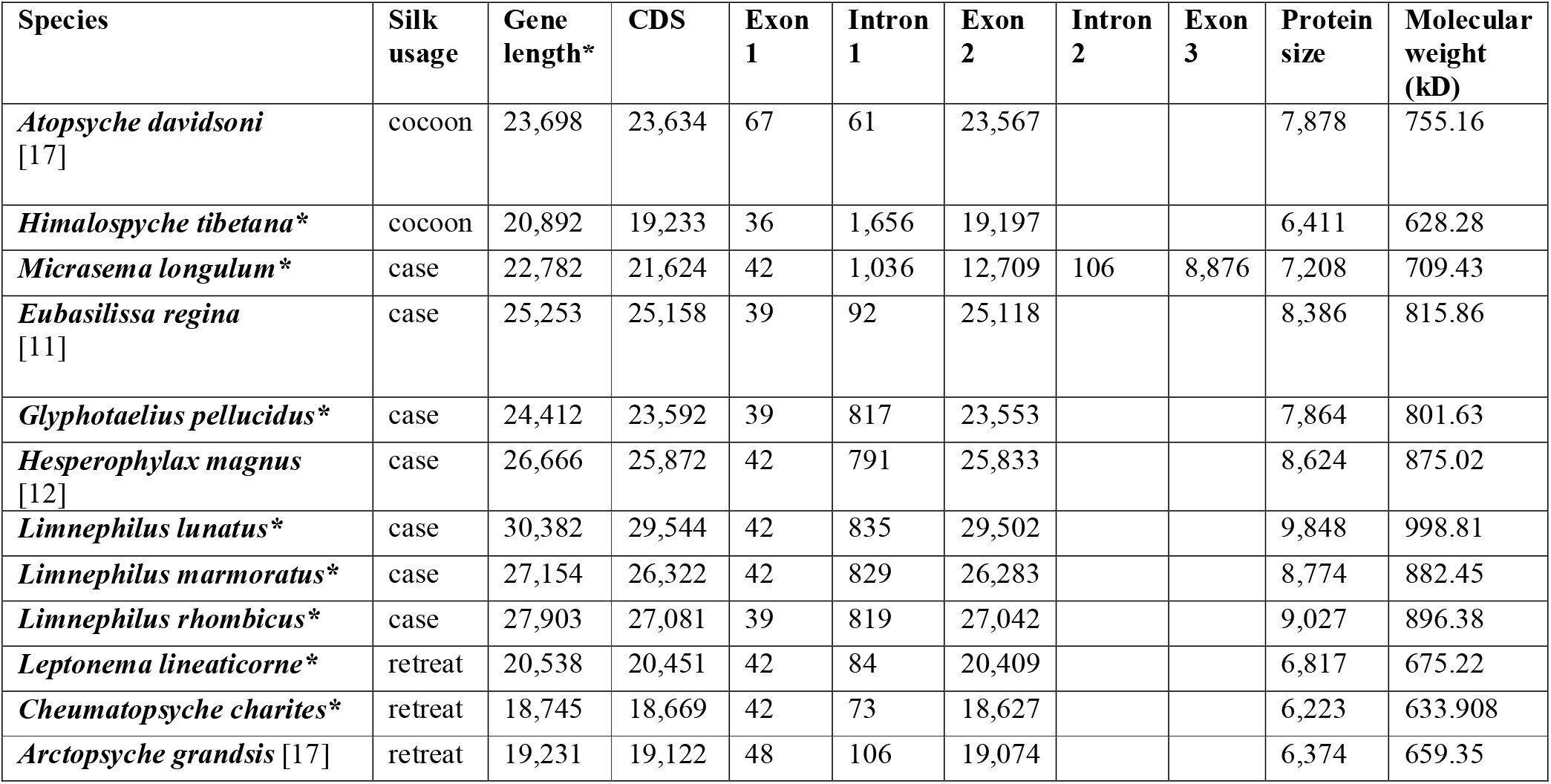
Full-length *h-fibroins* of twelve caddisfly species derived from long-read sequencing ordered by silk usage. Gene length including introns and stop codon; CDS= protein coding DNA, gene length, CDS, exon, intron in bp; protein size in amino acids. *: this study

## Results

### H-fibroin structure

The present study expands the number and diversity of known full-length *h-fibroin* sequences in caddisflies. Our final taxon sampling covers *h-fibroin* sequences of the main clades of Trichoptera with different silk usage: fixed retreat-making Annulipalpia (n=3), cocoon-making basal Integripalpia (n=2), and tube case-making Integripalpia (n=7).

The structure of the *h-fibroin* gene across the 12 species revealed a similar organization of introns and exons. The *h-fibroin* gene was characterized by a short exon (36-48 bp) followed by a single intron (61-1,656 bp) and a long second exon (12,709-29,502 bp) leading to a total length of 18,745-30,382 bp. The *M. longulum* sequence had an additional intron and exon (table 2).

The h-fibroin protein consists of non-repetitive n- and c-termini that were highly conserved across the different clades. The n-terminus contained 105-117 residues (without the signal peptide, figure 1A). There were 40.2% identical sites (% of columns in the alignment where all sequence are identical) and 74% pairwise identity (% of pairwise residues that are identical in the alignment, including gap versus non-gap residues, but excluding gap versus gap residues) among the species. The c-terminus consisted of 40 residues with 32.5% identical sites, 65.8% pairwise identity, and a conserved cysteine at position 19 in the c-terminus alignment (figure 1B).

**Figure 1:**
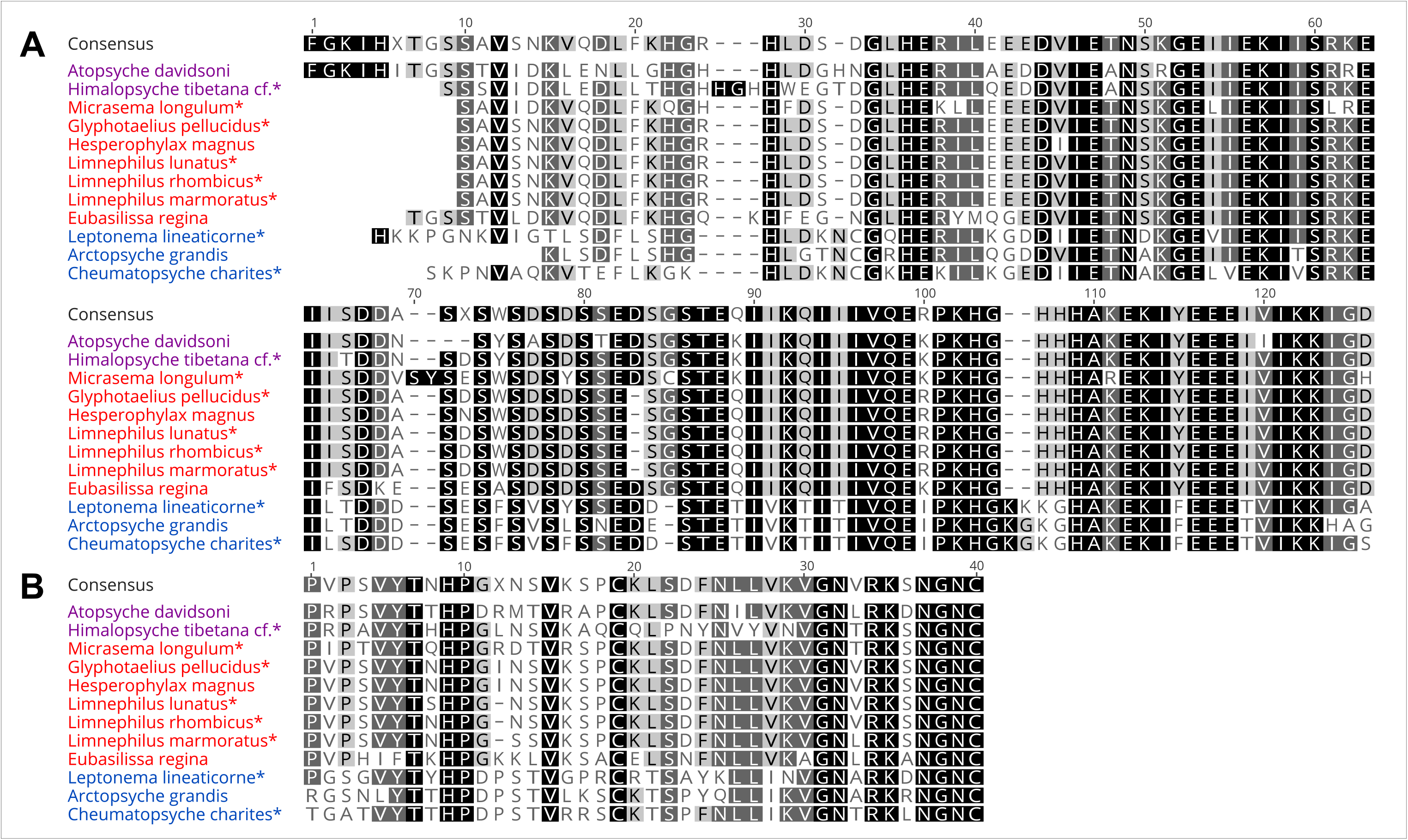
Highly conserved regions of the h-fibroin. A: n-terminus without signal peptide. B: c-terminus. Different silk usage is color-coded: violet: cocoon-, red: tube case-, blue: retreat-making.

The terminal domains flanked a central region, composed entirely of repeating sequence blocks, which have been represented in several ways in the literature to describe the primary structure of h-fibroin [4,8,20]. For example, Frandsen et al., [10] represented each unique (SX)nE motif as the beginning of a repeat. Structurally, this would correspond to defining each repeat as beginning with a single (SX)_n_E ß-strand [5]. In presenting the new h-fibroin sequences reported here, we have defined the repeating modules as each having two parts: first, a region comprising a variable number (1-6) of (SX)_n_E motifs, each separated by short (8-24) stretches of intervening amino acids, and, second, a variable length (8-144 residues) G(glycine) - or G(glycine)-P(proline)-rich region. Schematically, in figure 2, the two parts are represented by different symbols in each block. In the following paragraphs, we describe the repeating structural modules for each species analyzed in this study in detail. The full-length h-fibroin protein sequences are provided in supplementary notes 2-13. Schematic visualizations of each genus are presented in supplementary figures 5-14.

**Figure 2:**
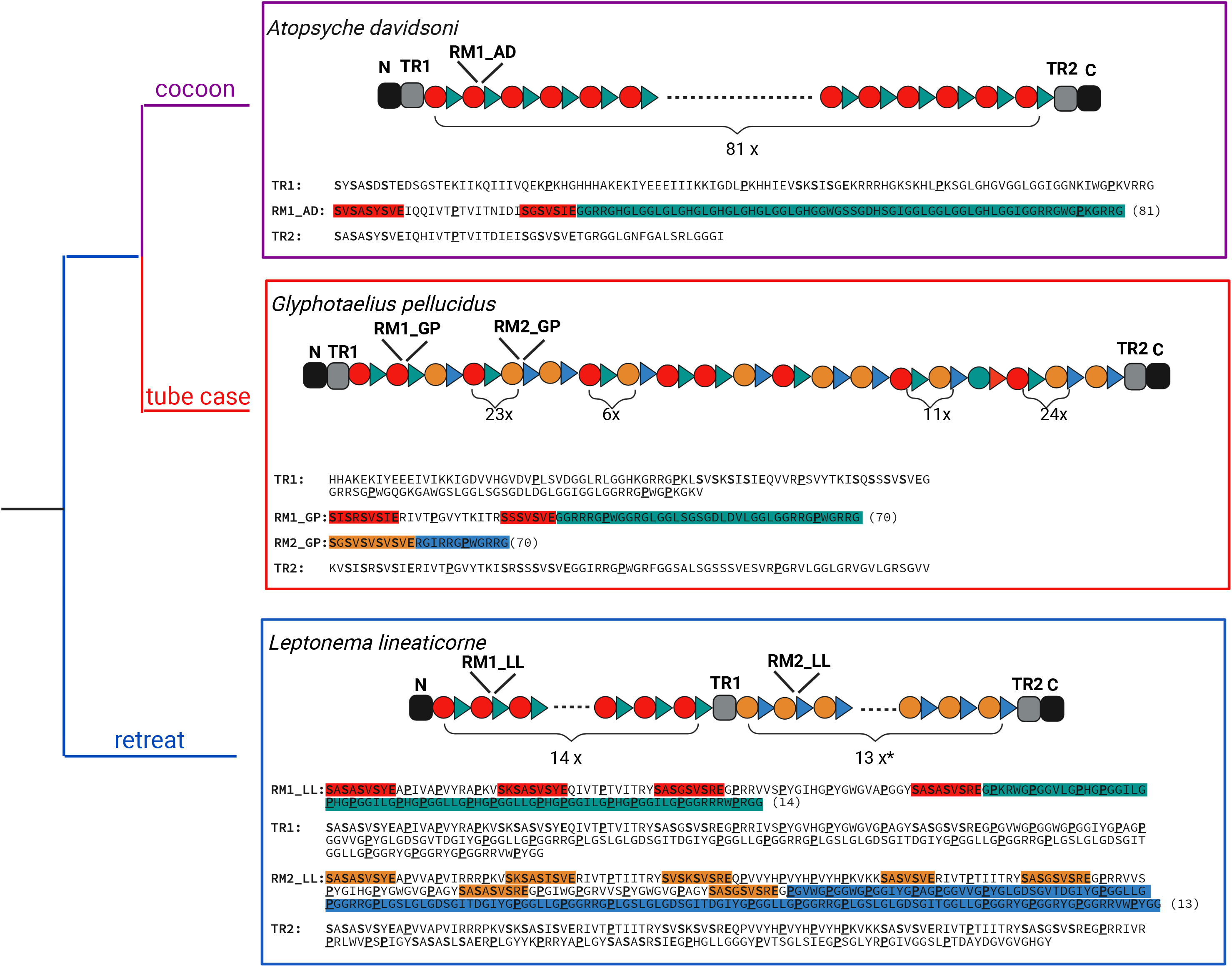
Schematic visualization of the primary structure of the h-fibroin gene of one representative species per clade. Black boxes: n- and c-terminus, for sequences see figure 1. Grey boxes: transition regions (TR). The repeating modules (RM) have two parts which are represented by different symbols: A region comprising a variable number (1-6) of (SX)_n_E motifs (red and orange circles correspond to red/orange residues in the sequence), each separated by short (8-24) stretches of intervening amino acids, and a variable length (8-144) G(glycine) - or G(glycine)-P(proline)-rich region (blue arrows correspond to blue residues in the sequence). Cocoon maker: *Atopsyche davidsoni* (AD): RM1_AD: Repeat module 1 occurs 81 times. Tube case maker: *Glyphothaelius pellicudulus* (GP): RM1_GP: Repeat module 1 and RM2_GP: Repeat module 2 occur 70 times each. Retreat maker: *Leptonema lineaticorne* (LL): RM1_LL: Repeat module 1 is repeated 14 times and RM2_LL: Repeat module 2 is repeated 13 times. The consensus sequence of each repeat module is given. The Glycine/Glycine-Proline-rich motifs vary in length. For full-length sequence see supplementary notes 2-13. Similar figures for each genus are given in supplementary figures 5-14. Phylogeny shown after [1]. Figure created with BioRender.com

### Basal Integripalpia (cocoon makers)

The structure of the repetitive central region in cocoon-building *A. davidsoni* was simple compared to the other sequences investigated in this study (figure 2, cocoon). The repeating module was embedded between two transition regions (TR), each of which occurs once and directly flanks the termini. In these transition regions, the sequence transitions from one type of module to another. The TRs had similarities to the repeating module but their sequence is unique, e.g. they occurred only one time in the h-fibroin and, in most cases, transition between the conserved termini and the repeating structural modules (figure 2: cocoon, tube case) or between repeat modules (figure 2: retreat). The h-fibroin of *A. davidsoni*, included a single repeating module consisting of a (SX)_4_E[15](SX)_3_E region and a G-rich region of variable length (40-70 residues). The number in the square bracket refers to the short stretches of intervening amino acids. This module repeats 81 times across the sequence.

The repetitive region of *Himalopsyche tibetana* was embedded between two transition regions. There were two repeating modules which are very similar. RM1 consists of a (SX)_6_E[16](SX)_4_**E**[15](SX)_6_E[58](SX)_5_E or (SX)_6_E[16](SX)_4_**D**[15](SX)_6_E[58](SX)_5_E (D: aspartic acid) motif and a glycine-rich motif of variable length (93-106 residues). RM2 was reduced to (SX)_6_E[16](SX)_4_E[15](SX)_6_E. The glycine-rich motif was 56 to 90 residues long. RM1 occurred 26 and RM2 occurred 13 times.

### Integripalpia (tube case makers)

The primary structure of the h-fibroin was variable within tube case-making Integripalpia. In all seven species sampled, the repetitive central region was flanked by two transition regions. However, the number and organization of repeat modules was diverse within this clade.

Specifically, in *E. regina* which uses plant material (leaves) and silk to build a tube case, the h-fibroin had only one type of repeating module consisting of a (SX)_4_E[8](SX)_4_E[17](SX)_4_E[11](SX)_3_E[20](SX)_3_E motif and a glycine-rich region (14-144 residues). It was repeated 39 times.

In *G. pellucidula* whose silk usage is similar to *E. regina*, the h-fibroin consisted of two repeat modules. RM1 consisted of a (SX)4E[13](SX)3E motif and glycine-rich region (38-81 residues), and RM2 consisted of a single (SX)_5_E motif and glycine-rich region (12-51 residues). Each RM was repeated 70.

Within the three *Limnephilus* species, the primary structure of the h-fibroin was comparable (supplementary notes 8-10). This genus uses a diverse array of plant materials (wood, moss, leaves) for tube case-making. There were two repeat modules. In *L. lunatus*, RM1 consisted of a (SX)_5_E[14](SX)_4_E[11](SX)_4_E region and a glycine-rich region (14-82). It occurred 96 times and was interrupted by ten RM2 which consisted of a single (SX)_5_E motif and a glycine-rich region (14), similar to *G. pellucidula* (figure 2). In *L. marmoratus*, RM1 occurred 68 and RM2 17 times. In *L. rhombicus* RM1 occured 59 and RM2 51 times.

The h-fibroin of *H. magnus*, which uses stones the build tube cases, was constituted by two repeat modules. RM1 contains a (SX)_5_E[14](SX)_4_E[11](SX)_4_E motif and a glycine-rich region (41-81 residues) and occurred 69 times and was interrupted by 13 RM2, which consisted of a single (SX)5E motif and a glycine-rich region (45 residues).

In the h-fibroin of *M. longulum* which constructs tube cases purely made of silk, there were three repeat modules. RM1 contained a (SX)_4_E[18](SX)_3_E[11](SX)_4_E[15](SX)_4_E motif and a glycine-rich region (41-58) and was repeated 24 times. RM2 consisted of a (SX)_4_E[18](SX)_3_E[11](SX)_4_E[18](SX)_3_E[11](SX)_4_E[14](SX)_4_E motif and a glycine-rich region (58 residues) which occurred 7 times. RM3 consisted of a (SX)_4_E motif and glycine-rich region (8-57) which occurred 36 times.

### Annulipalpia (fixed retreat makers)

In *Leptonema lineaticorne*, the sequence was divided into two parts each with a distinct repeat module (figure 2). RM1 consisted of a (SX)_4_E[13](SX)_4_E[11](SX)_4_E[24](SX)_4_E motif and glycine-rich region (68-92) and is repeated 14 times. RM2 consisted of a (SX)_4_E[14](SX)_4_E[11](SX)_4_E[19](SX)_3_E[11](SX)_4_E[24](SX)_4_E[23](SX)_4_E motif and a glycine-rich region (75-84 residues) and was repeated 13 times. There were two transition regions. TR1 in *L. lineaticorne* separated the two repeat modules and TR2 was located between RM2_LL and the c-terminus.

In *Cheumatopsyche charites*, the repetitive region was surrounded by two transition regions. Similar to *L. lineaticorne*, the h-fibroin was divided in two parts. The first part of the gene consisted of six repeat modules (RM1: SX_4_E[20]-SX_4_E and a glycine-proline-rich motif (112-141 residues): occurred 1x, RM2: (SX)_4_E[15](SX)_4_E[11](SX)_4_E[25]-(SX)_4_E[20](SX)_4_E and a glycine-proline-rich motif (86-118 residues): occurred 2x, RM3: (SX)_4_E[15](SX)_4_E[11](SX)_4_E and a glycine-proline-rich motif (44): occurred 1x, RM4: (SX)_4_E[15](SX)_4_E[11](SX)_4_E[25](SX)_4_E[20](SX)_4_E[20](SX)_4_E and a glycine-proline-rich motif (86-118): occurred 6x, RM5: (SX)_4_E[15](SX)_4_E[13](SX)_3_E[16](SX)_4_E[13](SX)_3_E[18](SX)_3_E[11](SX)_4_E and a glycine-proline-rich motif (89): occurred 2 times, RM6: (SX)_4_E[20](SX)_4_E[20](SX)_4_E and a G-P-rich motif (86): occurred 1x). In the second part of the RM1 and RM3 are alternating 13 times.

In *Arctopsyche grandis*, the sequence included three internally repeating modules flanked by two transition regions. RM1 consisted of a (SX)_3_E[12](SX)_4_E[12](SX)_4_E and a glycine-proline-rich motif (40-132 residues) and was repeated 24 times. RM2 contained a (SX)3E and a glycine-proline-rich motif (22-63). It occurred 31 times. RM3 comprised a (SX)_4_E[12](SX)_3_E and glycine-proline-rich motif (15-16 residues) and was represented 20 times.

### Amino acid composition

In general, the amino acid composition of the protein sequence was conserved across the taxa that we sampled. Glycine and serine were consistently the most abundant residues across all three clades. However, despite these consistent patterns in composition, we observed some differences among clades. In retreat-making caddisflies, h-fibroin was characterized by a high amount of proline which ranged from 9.9 to 12.3% (n=3). In contrast, the proportion of proline was much lower in the h-fibroin of tube case makers, ranging from 2.4 to 5.6% (n=7). In the h-fibroin of cocoon-making caddisflies, content of proline was even lower, ranging from 2.1-2.7% (n=2). When comparing the amino acid composition of h-fibroins in Trichoptera to those of terrestrial Lepidoptera, we find some consistent differences (table 3). While h-fibroins of both orders had high proportions of glycine (Trichoptera: 21.2-35.6%, Lepidoptera: 18.3-45.9%) and serine (Trichoptera: 9.3-17.2%, Lepidoptera:12.1-18.5%), h-fibroins in Lepidoptera had much more alanine (Trichoptera:0.1-4.9%, Lepidoptera: 21.9-30.3%). In addition, the Lepidoptera sequences exhibited a smaller percentage of charged residues. Negatively charged amino acids (aspartic acid and glutamic acid) ranged from 4-7.7% in the Trichoptera sequences but only 1.1-2.4% in Lepidoptera. Positively charged amino acids (arginine, lysine) summed up to 7.6-17.1% in Trichoptera h-fibroins, but were much lower in Lepidoptera (0.5-0.9%). Moreover, the amount of leucine and isoleucine was higher in h-fibroins of caddisflies compared to Lepidoptera. Specifically, the amount of leucine in the sequence ranged from 5.2% in retreat makers to 6.87% in case makers culminating in 10.1% in cocoon makers, whereas in Lepidoptera it was as low as 2.83%. The amount of isoleucine, rages from 4.77% (case makers), 5.2% (retreat makers) to 7.3% in cocoon makers. H-fibroins of Lepidoptera only contained 1.73% of isoleucine.

**Table 3:** Amino acid composition of full-length h-fibroins of Trichoptera and terrestrial sister order Lepidoptera. The mean is given for each amino acid and group of Trichoptera / Lepidoptera. Standard derivation is given in brackets. Amino acid composition for each individual is given in Additional data Files: DataS1-S17.

| Residue | Trichoptera -<br>retreat (n=3) | Trichoptera -<br>cocoon<br>(n= 2) | Trichoptera -<br>case<br>(n=7) | Terrestrial<br>Lepidoptera<br>(n=3) |
| --- | --- | --- | --- | --- |
| Gly (G) | 23.60 (1.71) | 33.55 (2.05) | 29.60 (1.66) | 31.05 (9.82) |
| Arg (R) | 9.43 (1.59) | 7.5 (1.5) | 12.17 (2.85) | 0.48 (0.11) |
| Lys (K) | 1.67 (0.48) | 1.3 (0.3) | 2.03 (0.53) | 0.23 (0.11) |
| His (H) | 2.03 (0.97) | 5.25 (1.75) | 1.16 (2.07) | 0.08 (0.04) |
| Ala (A) | 4.20 (0.50) | 1.75 (0.65) | 0.51 (0.49) | 26.13 (2.97) |
| Val (V) | 8.43 (0.37) | 6.25 (1.05) | 9.56 (0.83) | 3.18 (1.07) |
| Leu (L) | 5.20 (0.71) | 10.1 (0.7) | 6.87 (1.18) | 2.68 (2.35) |
| Ile (I) | 5.20 (0.22) | 7.3 (0.8) | 4.77 (1.08) | 1.53 (1.05) |
| Pro (P) | 10.77 (1.09) | 2.4 (0.3) | 4.44 (1.05) | 3.00 (1.91) |
| Ser (S) | 11.77 (0.50) | 12.1 (2.8) | 15.37 (1.25) | 15.35 (2.32) |
| Tyr (Y) | 5.10 (1.36) | 2.45 (1.35) | 3.70 (2.08) | 4.75 (1.34) |
| Asp (D) | 3.37 (1.39) | 1.5 (0.3) | 2.49 (0.73) | 0.55 (0.17) |
| Glu (E) | 2.60 (0.14) | 2.5 (0.1) | 3.04 (0.25) | 1.35 (0.63) |
| Thr (T) | 2.90 (0.37) | 3 (0.2) | 1.49 (0.43) | 0.93 (0.43) |
| Trp (W) | 1.87 (0.57) | 1.35 (0.25) | 2.33 (0.88) | 0.05 (0.09) |

## Discussion

In this study, we report eight new full-length *h-fibroin* sequence for caddisflies across a diverse set of silk usage. Three of these were generated from new genomic resources, while five were mined from previously published genomes, highlighting the relevance of genome sequencing projects for the wider scientific community. The new full-length *h-fibroins* represent a substantial increase in the number of genomic resources available for the study of caddisfly silk and allowed us to compare the major silk gene for twelve species across the primary clades of Trichoptera, in which species exhibit different silk usage.

The genetic structure of *h-fibroin* of the twelve Trichoptera species showed similar organization of introns and exons, in line with previously reported *h-fibroin* sequences of Lepidoptera [11,21]. In addition to resolving the genetic structure, we compared the primary protein sequence of the h-fibroins. Consistent with previous research [4,10], we found that structural elements of h-fibroin are conserved across Trichoptera. For example, of the species sampled, the n- and c-termini exhibit a high pairwise identity (n-terminus: 74%, c-terminus: 65.8%). In Lepidoptera the conservation of the termini is linked with function. The n-terminus dimerizes and the c-terminus has been reported to interact with the light chain fibroin: the terminal cysteine forms an intermolecular disulfide bond with the light chain fibroin in the silkworm *Bombyx mori* [22]. Given similar patterns of conservation of the termini in Trichoptera, their function is likely conserved across both orders. The c-terminus of all Trichoptera was characterized by the presence of a cysteine at position 19 in the alignment (figure 1). This suggests that disulfide crosslinking of fibroins occurs also in caddisflies and implies that covalent complex formation though the c-terminus is of similar importance for the structure, stability, and secretion of the fibroin complex as reported in *B. mori* [23]. In addition to conservation of the termini, some conserved themes emerged in the central repetitive region, which consisted of repeating two-part structural modules, each containing a characteristic region of (SX)_*n*_E motifs interspersed with glycine-rich (in cocoon- and case makers) or glycine-proline-rich (in retreat makers) regions of variable length. The serines of the (SX)*n*E motifs have been shown to be extensively phosphorylated [6–8]. These phosphates then bind multivalent metal ions, which stabilize the silk and are responsible for the strength of the silk [5,24,25]. A structural model has been proposed, in which each (pSX)nE motif forms a ß-strand, which in turn associates into anti-parallel Ca2+-stabilized ß-sheets [5]. The ß-sheets stack through alternating hydrophobic and Ca2+-phosphate interfaces creating microcrystalline β-domains. By this model, each repeating module would correspond to a structure in which anti-parallel ß-sheets are linked with a glycine-rich or glycine-proline-rich spacer region. The (SX)_n_E/glycine-rich blocks may combine to form a higher-order ß-domain structure through intra- or intermolecular stacking of the [(SX)_n_E]_m_ ß-sheets. The crystalline ß-domains would be separated by flexible and extensible glycine-/glycine-proline-rich regions. These signatures of conservation in repeat modules despite ~280 million years of divergence [1] and diverse silk usage, suggest a common mechanism for protein folding and silk formation across the underwater silks generated by all clades as suggested by [10].

Despite the conservation in structure, we observed variation in the ordering and number of repeat modules. Our study builds on previous studies (i.e.[4,20,5,8,10,12,17]). by unveiling substantial diversity in the number and order of these repetitive structures. We observed a range of complexity in the repetitive structures across the phylogeny. In the simplest h-fibroin structure, sequenced from *A. davidsoni*, a single structural module was repeated 81 times (figure 2). Slightly more complicated was the h-fibroin sequence of tube case-making *G. pellucidula*, which consisted of two repeat modules shuffled throughout the central region of the sequence (figure 2). The sequences of the fixed retreat-makers were more variable, including *L. lineaticorne*, which was split into two parts, each of which consisted of a unique repeating module (figure 2). The variations in repeating block patterns between the different clades may reflect adaptations for the diverse silk usage. For instance, cocoon-making *Himalopsyche tibetana* and *Atopsyche davidsoni* are free-living as larvae and only produce pupal silk for building cocoons and pupal domes. The increase in h-fibroin sequence complexity scaled with silk usage diversity. The species in this study that construct both fixed retreats and capture nets exhibited the most variable h-fibroin sequences. The diversity in the number and order of the repeat modules may hold clues to unraveling the unique applications of silk across clades (e.g., case-, cocoon-making vs. fixed retreat-building) and may be directly responsible for differences in mechanical properties. Amino acid composition of the h-fibroins was largely conserved across samples, with some notable exceptions. The proportion of proline was clade specific. While proline was found in low proportions in the h-fibroin of Integripalpia (cocoon (2.1-2.7%) and tube case makers (2.4-5.6%)), higher proportions were found in Annulipalpia sequences (9.9-12.3%). Annulipalpian larvae are generally characterized by their fixed retreats which serve as shelters. In addition, some annulipalpian families, including all of the species investigated in this study, also construct silken capture nets, which are used for capturing food and would presumably require more extensible silk. Future work in caddisflies, should focus on linking physical properties of the silk with variation in h-fibroin sequence. For example, orb-weaver spiders use silk to build prey capture spirals with high fiber extensibility. This is necessary to catch insects in flight without breaking the web. The abundance of proline content in the major ampullate spidroin MaSp2 of orb-weaver spider silk was linked to enhanced extensibility of these fibers [26–31] because it increases the secondary structure disorder in the amorphous region [26,28,27]. Furthermore, Arakawa et al. [32] found that breaking strain was positively correlated the presence of an amorphous, proline-rich region in the major ampullate spidroins MaSP1/2, which are often incorporated into dragline threads [32].

Despite similarities in the gene structure of *h-fibroin* in aquatic Trichoptera and terrestrial Lepidoptera, we found consistent differences in amino acid composition of the protein sequences. In general, to prevent limitation by protein content in their diets, non-essential amino acids glycine, alanine, and serine are the dominant residues in insect silk genes [33]. However, in contrast with Lepidoptera h-fibroins, which have high alanine content, Trichoptera h-fibroin sequences were extremely low in alanine. This is likely due to differences in how the proteins are folded. In Lepidoptera, the ß-sheet structures are mainly derived from repetitive polyglycine-alanine (*B. mori*) and (non-)/polyalanine-domains (saturniid moths) and thus these amino acids are important for the strength of the silk [34,35]. In contrast, as noted above, caddisfly h-fibroin ß-sheets are primarily formed through the interaction of phosphorylated serine blocks with metal ions derived from their aquatic environment, an adaptation specific to aquatic dwelling species [25]. H-fibroins of both, Trichoptera (9.3-17.2%) and Lepidoptera (12.1-18.5%), contain high % of serines. Therefore, Ashton et al. [3] suggested that one of the key molecular adaptations of a terrestrial ancestor silk to aquatic environments could have been kinase phosphorylating H-fibroin serines. We also detected a much higher percentage of charged residues in caddisflies (table 3), which has previously been hypothesized as an adaptation for aquatic silks [4].

### Potential implications

The new genomic data provided in this study was used to investigate the primary structure of h-fibroins across caddisflies and thus presents further insights into the genomic basis of adhesive underwater silk in Trichoptera. While we observed conserved patterns in the primary structure of the h-fibroin, the amino acid composition and the number and arrangement of repeating modules varied among species with different silk usages. The next step toward a better understanding of the role of this variation in generating the myriad silk phenotypes that we observed across Trichoptera is to link these sequences with experimental evidence, such as mechanical testing as shown for spider silk [32]. Such studies are essential to gauge the potential of caddisfly silk in material science, and the sequences that we have compiled here represent an important step toward performing such analyses.

## Methods

We collected a single larva of *M. longulum* in Germany, Hesse, Bieber (5009’39.7’N, 920’09.1’’E), a single adult individual of *L. lineaticorne* in an Amazon Blackwater channel in Ecuador, Sucumbios, Sacha Lodge (Amazon Basin), Caño Anaconda (0°28’20.71’’S; 76°27’59.08’’W, elevation 237 m asl.) and a single larva of *H. tibetana* in Gasa, Bhutan (28°03.9477’N, 89°39.0019’E, elevation 3824 m asl.). We extracted high molecular weight from single individuals and prepared DNA sequencing libraries following the instructions of the SMRTbell Express Prep. For each individual, one SMRT cell sequencing run was performed on the Sequel System II in CCS (circular consensus sequencing) mode using 30-hour movie time. For *M. longulum*, we generated genomic HiFi reads by CCS from the subreads.bam files provided by the sequencing facility using pbccs tool v6.6.0 in the pbbioconda package (https://github.com/PacificBiosciences/pbbioconda). For *L. lineaticorne* and *H. tibetana* we generated HiFi reads from the raw data (reads with quality above Q20) using PacBio SMRTlink software (https://github.com/PacificBiosciences/pbbioconda). We estimated genome size using sequencing reads and a *k-mer*-based statistical approach. After counting *k-mers* with JELLYFISH v2.2.10 [36] using jellyfish count -C -s 25556999998 -F 3 and a *k-mer* length of 21 (-m 21) with the ccs-reads, we produced a histogram of *k-mer* frequencies with jellyfish histo. We ran GenomeScope 2.0 [37] with the exported *k-mer* count histogram within the online web tool (http://qb.cshl.edu/genomescope/genomescope2.0/) using the following parameters: *k-mer* length = 21, Read length = 150, Max kmer coverage = 1000. We assembled the *L. lineaticorne* and *H. tibetana* genome, with hifiasm [38] with the default settings. Contaminations of adapters detected by NCBI in the *L. lineaticorne* genome assembly were filtered out with samtools faidx [39]. We used flye v2.8.2 [40] with the –pacbio-hifi option to assemble the *M. longulum* genome. We evaluated the assembly quality based on continuity (QUAST v5.0.2 [41]) and completeness of Benchmarking Universal Single-Copy Orthologs (BUSCOs) with BUSCO v5.2.2 [42,43] using the lineage dataset endopterygota_odb10 in genome mode. In addition, we calculated the back-mapping-rate of the HiFi reads to the assemblies using backmap.pl (Pfenninger et al. 2022; Schell et al. 2017) with the parameter -hifi. Other parameters were kept as default. This wrapper script automatically maps the reads to the assembly with minimap2 [44] and executes qualimap [45], MultiQC [45], bedtools [46], and RScript (R Core Team 2021) to create the mapping quality report and a coverage histogram. In additions, it plots the coverage distribution and estimates of genome size from mapped nucleotides divided by mode of the coverage distribution (>0). The final genome assemblies were screened for potential contaminations with taxon-annotated GC-coverage (TAGC) plots using BlobTools v1.1.1 [19]. For this purpose, the bam file resulting from the backmapping analysis was converted to a blobtools readable cov file with blobtools map2cov. Taxonomic assignment for BlobTools was done with blastn 2.10.0+ [47] using -task megablast and -e-value 1e-25. The blobDB was created and plotted from the cov file and blast hits.

Recently, the Wellcome Sanger Institute published four high-quality caddisfly genomes (*Glyphotaelius pellucidus:* GCA_936435175.1, *Limnephilus lunatus:* GCA_917563855.2, *Limnephilus marmoratus*: GCA_917880885.1, *Limnephilus rhombicus:* GCA_929108145.1). An additional high-quality genome was published by [48]. We identified the *h-fibroin* genes in these assemblies by using tBLASTn to search the assemblies with the conserved n- and c-termini with query sequences from previously published species *Hesperophylax sp*. [5], *Limnephilus decipiens* AB214509 [4] and *Rhyacophila obliterata* AB354689.1 and AB354588.1 [13] in Geneious Prime 2022.1.1 (https://www.geneious.com). After verifying that both BLAST hits (hit with n- and hit with c-terminus) were isolated to the same contig in the genome assembly, we extracted the sequences and 1,000 bp of flanking regions from the assembly using the sequence view “extract” in Geneious and annotated this region using Augustus v.3.3.3. [49]. Introns that did not affect reading frames were manually removed from the annotation and h-fibroins were manually curated as described in supplementary note 14. Protein coding nucleotide sequences were translated with the Geneious Tool “Translate” using the standard genetic code. We used the same approach to extract and annotate the *h-fibroin* in *L. lineaticorne* but using the termini of *Parapsyche elsis* [10]. For *H. tibetana* we used termini of *Rhyacophila obliterata*. For details see supplementary table 1.

Because the sequencing coverage and thus the assembly quality was low for *M. longulum*, we had to conduct several extra steps to identify the *h-fibroin* in this species. With query sequences from the termini of *Hesperophylax sp*., we used tBLASTn to identify the conserved n- and c-termini and extracted the *h-fibroin* plus 500 bp flanking regions from the flye assembly as described above. Recently, researches have shown that there is substantial haplotypic variation within individuals for genes underlying the primary silk proteins in both insects and spiders [17] and the silk gene we extracted of the *M. longulum* assembly was probably not haplotype resolved and thus riddled with frameshift errors and stop codons. To assemble the error-free full-length sequence of the *h-fibroin*, we mapped all CCS reads back to the extracted region of the genome and used FreeBayes v1.3.5 [50], a haplotype-based variant detector, to call variants and WhatsHap v.1.1 phase [51] to phase genomic variants. To reconstruct the primary haplotype from the phased VCF file, we used the bcftools consensus command. We annotated the resulting sequence with Augustus v.3.3.3 as described above (details given in supplementary note 14).

We predicted signal peptides and the location of their cleavage site of the h-fibroin protein sequences with the SignalP 6.0 server (https://services.healthtech.dtu.dk/service.php?SignalP,[52]) using the following settings: organism = Eukarya, model mode = slow.

### Comparison of heavy-chain fibroins across caddisflies clades

We used the previously published full-length h-fibroin sequences and the newly identified h-fibroin sequences generated here to compare their primary structure. In order to compare conserved regions of the h-fibroin proteins, we aligned the n- and c-terminus each (without the signal peptide) in Geneious using the Muscle 3.8.425 [53] plugin with a maximum of 1,000 iterations. We compared % pairwise identity and % of identical sites in Geneious. For each species, we used custom-made scripts to split the silk gene into repeat modules (https://github.com/AshlynPowell/silk-gene-visualization/tree/main). For schematic visualization of the primary structure, we generated a consensus sequence of all representative sequences of each repeat module by aligning these in Geneious using the Muscle 3.8.425 plugin with a maximum of 1,000 iterations. We used Expasy ProtParam (https://web.expasy.org/protparam/) to compute the molecular weight and the amino acid composition of each sequence. For comparison, we also calculated amino acid composition of the four available Lepidoptera full-length h-fibroins [(silkworm *Bombyx mori*, Zhou et al, 2000; Indianmeal moth *Plodia interpunctella*,[11]; painted lady *butterfly Vanessa carduii* [17] and spindle ermine moth *Yponomeuta cagnagella* [21]] using ProtParam.

## Author contributions

J. Heckenhauer: Conceptualization, Data curation, Formal analysis, Investigation, Validation; Visualization, Writing - original draft; Writing - review & editing

R. J. Stewart: Conceptualization, Investigation, Validation; Writing - original draft; Writing - review & editing

P. B. Frandsen: Conceptualization, Formal analysis, Funding acquisition, Investigation, Resources: computational, Writing - original draft; Writing - review & editing

B. Ríos-Touma: Resources: material for DNA extraction, Writing - review

T. Dorji: Resources: material for DNA extraction, Writing - review

A. Powell: Software; Writing - review

S. U. Pauls: Conceptualization, Funding acquisition, Project administration; Resources: computational; Writing - review & editing

## Acknowledgements

We thank Brigham Young University and LOEWE-Centre for Translational Biodiversity Genomics (TBG) for providing the computational resources needed to complete this study.

The *Leptonema lineaticorne* specimen was collected in Ecuador under the “Genetic Resources Access Contract” No. MAAE-DBI-CM-2021-0161, and the support of project AMB.BRT. 19.02 (“Mountain Freshwater Diversity, from Taxonomy to Functional Genomics, and Approximation from Trichoptera”). We appreciate help from Sacha Lodge for field logistics.

## Availability of source code

All custom-made scripts used in this study are available on GitHub.

Project name: h-fibroin-visual

Project home page: https://github.com/AshlynPowell/h-fibroin-visual

## Data Availability

Raw sequence data, genome assembly and sample information has been submitted to NCBI.

*L. lineaticorne:* Raw sequence data: SRR20711493, genome assembly: JANIJI000000000, sample information: SAMN25408291). *M. longulum:* Raw sequence data: SRR15840268, sample information: SAMN20982427. *H. tibetana:* Raw sequence data: SRR22537910, sample information: SAMN31697436, genome assembly: JAPJYX000000000. The data supporting the results of this article (assemblies, all data associated with quality control of the assemblies, and full-length *h-fibroin* gene sequences including introns, *h-fibroin* protein coding nucleotide sequences and h-fibroin protein sequences newly identified in this paper) is available in the Figshare repository: https://figshare.com/s/03f88091eda258465d2b.

